# GWAS meta-analysis of resistance against Piscirickettsia salmonis in Atlantic salmon

**DOI:** 10.1101/2022.12.24.521873

**Authors:** Marín-Nahuelpi Rodrigo, Baltasar F. Garcia, Agustin Piña-Elgueda, Jousepth Gallardo-Garrido, Paulina López, Daniela Cichero, Thomas Moen, Jørgen Ødegård, José M. Yáñez

**Author notes:** Corresponding author: Facultad de Ciencias Agronómicas, Universidad de Chile, Avenida Santa Rosa 11735, 8820808, La Pintana, Santiago, Chile.

## Abstract

Salmonid rickettsial syndrome (SRS) remains as one of the most important pathogens for salmon farming. Genetic improvement has proven to be a viable alternative to reduce mortality in breeding stock. Understanding the genetic architecture of resistance has been a matter of ongoing research aimed at establishing the most appropriate method by which genomic information can be incorporated into breeding programs. However, the genetic architecture of complex traits such as SRS resistance may vary due to genetic and environmental background. In this work, we used the genotypes of a total of 5839 Atlantic salmon from 4 different experimental challenges against *Piscirickttsia salmonis*, which were imputed high density (∼930K SNP) to perform within-population genomic-association analyses, followed by a meta-analysis of resistance to SRS defined as binary survival and day of death. The objectives of this study were to i) uncover the genomic regions associated with resistance to SRS among multiple populations; and ii) identify candidate genes associated with each trait definition. SNP-based meta-analysis revealed a clear QTL on *Ssa02* for both traits while gene-based meta-analysis revealed 16 genes in common for both traits. Our results suggest a polygenic genetic architecture and provide novel insights into the candidate genes underpinning resistance to *P. salmonis* in *Salmo salar*.

## INTRODUCTION

*Piscirickettsia salmonis* is a facultative intracellular bacterium causing the infectious disease salmonid rickettsial syndrome (SRS), being considered as one of the main pathogens threatening Chilean salmon farming (Rozas & Enríquez, 2014). Genetic improvement for resistance against infectious diseases is a feasible and sustainable alternative to reduce outbreaks in the field (Bishop & Woolliams, 2014). The evidence for genetic variation in resistance, defined as binary survival (BS) or day of death (DD), have already demonstrated the feasibility of implementing a breeding program in different aquaculture species (Barria et al., 2019; Bassini et al., 2019; Yáñez, Bangera, et al., 2016; Yáñez et al., 2013, 2014)

The advancement of molecular technologies has allowed the development of dense SNP-panels for salmonid species (Houston et al., 2014; Palti et al., 2015; Yáñez, et al., 2016), which have been useful in genomic prediction, and to explore the genetic architecture and potential genomic variants involved in the resistance against SRS in different salmonid species (Bangera et al., 2017; Barria et al., 2019; Correa et al., 2015; Moraleda et al., 2021; Yáñez et al., 2019; Yoshida et al., 2018). These studies have shown that SRS is a highly polygenic trait influenced by a large number of small effect associated variants. However, the genomic architecture of a trait may vary across populations and datasets, by the influence of different factors such as sample size, SNP density or genetic background, which may contribute to identifying spurious association signals or might not be enough to achieve a significant statistical power to correctly detect the significance and magnitude of associations (Sul et al., 2018).

Different methodologies have been developed over the last decades to address these shortcomings. Genotype imputation (Browning et al., 2018; Sargolzaei et al., 2014) has proven to be a useful tool for decreasing genotyping costs and increasing the accuracy of estimating genomic breeding values in salmonids (Tsai et al., 2017; Yoshida et al., 2018). In addition, it can also boost the power of GWAS due to an increased number of SNPs and the increased linkage disequilibrium (LD) between markers and causative variants (Das et al., 2018). GWAS meta-analyses is a common approach used to improve the power of detecting associations between genomic variants and traits of interest (Lin & Zeng, 2009). By combining the results of different studies together, meta-analysis tools allow detection of common genomic regions or genes affecting complex genetic diseases and traits of interest by improving the statistical power and accuracy of estimates (de Bakker et al., 2008; Sharifi et al., 2018)

Meta-analyses at individual-variant level (SNP-based) or gene-based level, aiming at evaluating chromosomal regions and genes involved in *P. salmonis* resistance across different cohorts, would help to provide further insights into *P. salmonis* resistance in Atlantic salmon. In the current study, we utilize large-scale genotype data and statistical genetic tools, to finely map genomic regions that underlie the genomic architecture of *P. salmonis* resistance. The objective of this study were i) detect the chromosomal regions involved in *Piscirickettsia salmonis* resistance using an individual SNP-based meta-analysis and ii) identify genes associated to *Piscirickettsia salmonis* resistance using a gene-based meta-analysis.

## MATERIALS AND METHODS

### Atlantic salmon samples and phenotypes

The populations used in this study were Atlantic salmon (*Salmo salar*) from three different genetic lines run in parallel in AquaGen Chile’s breeding program (Puerto Varas, Chile). This program was initially established using a diverse and representative collection of Atlantic salmon genetic material from 40 different Norwegian rivers during 1971 and 1974. These populations have been selected for several traits including growth-related, resistance against diseases (ISA, IPN, Piscirickettsiosis, Furunculosis), avoidance of early maturation, stress tolerance, skeletal deformities, and other traits for several generations. For a detailed description about rearing conditions and population management see AquaGen website (www.aquagen.cl).

Fish from 4 different experimental challenges performed at different years (POP A – POP B – POP C – POP D) belonging to 3 genetic lines were used in this study. A brief description of each challenge including genetic line and cohort, experimental design, time, strain, mortality, water type and average weights (if available) is shown in **Table 1**. A total of 5,858 animals were challenged and genotyped across all cohorts. Fish were intraperitoneally (IP) injected with a lethal dose (LD_50_) of different strains of *P. salmonis* inoculum or exposed by cohabitation (COH) with IP fishes using fresh or saltwater. Resistance to *P. salmonis* was defined as: 1) time to death (TD), measured in days, with values ranging from 1 until the end of the challenge test; and 2) binary survival (BS), with a value of 1 or 0 if the fish died or survived, respectively, until the end of the challenge.

**Table 1.** Description of the populations and challenges available for meta-analyses.

| POP<sup>1</sup> | N<sup>2</sup> | IW<sup>3</sup> | FW<sup>4</sup> | WT<sup>5</sup> | ED<sup>6</sup> | ST<sup>7</sup> | MOR<sup>8</sup> | DC<sup>9</sup> |
| --- | --- | --- | --- | --- | --- | --- | --- | --- |
| POP A (L1) | 1334 | - | 426.8 | SW | COH. | EM90 - A | 51% | 76 days |
| POP B (L1) | 1349 | - | 221.18 | FW | COH. | EM90 - A | 43% | 63 days |
| POP C (L2) | 1140 | 50.18 | - | SW | COH | EM90 - A | 39% | 67 days |
| POP D (L3) | 2016 | 42.8 | - | FW | COH. | EM90 - A | 50% | 69 days |

<sup>1</sup> Population (challenge); <sup>2</sup> Number of genotyped individuals; <sup>3</sup> Initial weight mean; <sup>4</sup> Final weight mean; <sup>5</sup> Water type; <sup>6</sup> Experimental design; <sup>7</sup> SRS strain; <sup>8</sup> Cumulative mortality; <sup>9</sup> Days of challenge.

Challenges were performed in VESO Chile Colaco (VCC) center. Before fish transport to facilities, a random sample (15-20) from each cohort were selected to evaluate the sanitary status of the population, i.e., by qRT-PCR for Infectious Salmon Anemia virus (ISAv), Infectious Pancreatic Necrosis virus (IPNv), Piscine orthoreovirus (PRV), *Piscirickettsia salmonis*, and *Renibacterium salmoninarum, Flavobacterium psychrophylum* and *Tenacibaculum dicentrarchi*. After tagging, fish were maintained in acclimatation tanks from 3 to 20 days prior to experimental challenges. Environmental parameters were measured throughout the challenge and the experimental challenge continued until the mortality curve showed a plateau. Daily mortality was recorded, and body weight was measured for each fish at the start of the challenge (initial weight = IW) and/or at time of death (final weight = FW). Surviving fish were euthanized and body weight was also recorded. Fin clips from all fish were sampled and stored in 95% ethanol at -80°C until they were genotyped.

### Genotyping and imputation

Populations were genotyped using a custom 65K SNP Affymetrix array developed for the AquaGen strain of salmon. A quality control was performed independently for each population. SNPs with call-rate < 0.9, minor allele frequency (MAF) < 0.01 and departing from Hardy-Weinberg equilibrium Bonferroni-corrected (*p-value* = 0.05 / N° SNPs remaining after previous filters).

For imputation we used two reference Atlantic salmon populations consisting of 1480 and 1314 animals genotyped with 200K and 930K, respectively. Both 65K and 200K SNP panels belong to a subset of the 930K XHD Ssal array (dbSNP accession numbers ss1867919552– ss1868858426) that is described in Barson et al., (2015). A stepwise imputation strategy was conducted, imputing first from 65K to 200K and then to 930K as this strategy offers higher accuracy of imputation (van Binsbergen et al., 2014). We performed a validation study to remove SNPs with low imputation accuracy using a five-fold cross validation scheme for both 200K and 930K datasets. For the 200K SNP validation study, 296 animals (20% of the total) were randomly assigned as reference and the remaining 1184 (80% of the total) were assigned as validation groups across the five validation groups. The reference animals had 209,579 SNPs and the validation individuals had their genotypes masked keeping only 43,766 SNPs that were in common among the four genotyped populations (“low density”). Imputation was performed for each validation group and the accuracy of genotype imputation was estimated using Pearson’s correlation (r²) between imputed and observed genotypes. We finally selected only SNPs with mean accuracy of imputation (r²) greater than 0.8 as the final set of imputable SNPs.

For the 930K validation (1314), the same strategy was implemented using 20% of animals with 633,254 SNPs and 80% of animals with 200,394 SNPs as reference and validation, respectively. All imputations were performed using the FImpute v3 software (Sargolzaei et al., 2014). All SNPs used in this work were positioned based on the penultimate Atlantic salmon reference genome (assembly ICSASG_v2) (Lien et al., 2016).

### Genome-wide association analyses

Four genome-wide association analyses (GWAS) were performed independently. GWAS analyses were performed using the MLMA-LOCO algorithm of GCTA software v1.93.2beta (J. Yang et al., 2011) which uses a genetic relationship matrix (GRM) to account for population structure and genomic relationships and the Leave One Chromosome Out (LOCO) method, which shows an increased analysis power when the SNP under exam is not itself associated with the phenotype but linked to causal SNP and avoids proximal contamination (Cheng & Palmer, 2013; J. Yang et al., 2014). The association study can be represented using the following model:

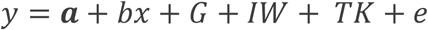

where *y* corresponds to the phenotype, *a* the mean value, *b* the fixed regression slope (beta) corresponding to the allele substitution effect of the SNP tested, *x* the SNP genotype (with levels 0/1/2 corresponding to the number of minor alleles in the genotype of an individual), *G* is the random genetic effect captured by the GRM, *IW* is the initial weight used as covariable if available, *TK* is the tank if the challenge was performed using different tanks and *e* is the residual random error.

To identify trait-associated regions, the statistical significance of the effect of each SNP was corrected for a genomic-level significance threshold according to the Bonferroni correction (*p-value* = 0.05/ total number of markers). Chromosome-level threshold significance was defined as *p-value* = 0.05/ total number of markers divided by the number of chromosomes. SNP-based heritabilities (h^2^_SNP_) were also estimated with GCTA using the REML method. Results were visualized as Manhattan plots using custom R scripts (v 4.1.3) (R Core Team 2022).

### Meta-analyses for genomic architecture

The meta-analysis for each trait was performed using METAL software (Willer et al., 2010). This software uses the weighted Z-scores model considering the *p*-value, direction of effects and weights the SNP significance depending on their presence within each individual GWAS input. Results were presented as Manhattan plots of the GWAS meta-analysis. Significance thresholds were calculated using the same criteria as individual GWAS analyses thresholds, i.e., genome-wide significance threshold using Bonferroni correction (*p*-value = 0.05/ total number of markers) and chromosome-level as *p-value* = 0.05/ total number of markers divided by the number of chromosomes.

### Meta-analyses for candidate genes

To identify candidate genes associated with each and both traits we performed a genome-wide gene-based meta-analysis using MAGMA (v1.10) (de Leeuw et al., 2015). This analysis uses the P-values from each variant-based GWAS analysis as input and tests the joint signal of all variants in a gene with each phenotype while accounting for LD between variants. SNPs were mapped to genes using ICSASG_v2 assembly. MAGMA gene-based meta-analysis uses a multiple linear PC regression model, and computes p-values for each gene using an F-test. We considered genes to be genome-wide significantly associated if its p-value was lower than a threshold value calculated using Bonferroni correction (0.05 / number of genes tested: *p*-value < 1.49 × 10^−06^). Additionally, we performed a gene-based enrichment analysis using the binary cut method implemented in the *simplifyEnrichment* package of R (Gu & Hübschmann, 2021). This method allows for clustering similar functional terms into groups of biological processes.

## RESULTS

### Descriptive statistics, imputation and heritabilities

Summary statistics for phenotypes (BS and TD) representing resistance to *P. salmonis* are shown in Table 2. Briefly, the longest challenge lasted 76 days with an average day-death of 64 days, and the shortest challenge lasted 60 days with an average day-death of 39 days. First death occurred at day 1, 23, 36 and 26 in POP A, B, C and D, respectively. Cumulative mortality at the end of each challenge ranged from 39% to 51%.

**Table 2.** Summary statistics for time to death (TD), binary survival (BS) measured in all populations.

| Trait | POP | Mean | SD | CV (%) | Min | Max | $\sigma^2_a$ | $\sigma^2_e$ | 930K SNP-h<sup>2</sup> |
| --- | --- | --- | --- | --- | --- | --- | --- | --- | --- |
| BS<br>(Status) | A | 0.51 | 0.50 | 98.03 | 0 | 1 | 0.06 | 0.18 | 0.25 ± 0.03 |
|  | B | 0.43 | 0.49 | 114.12 | 0 | 1 | 0.08 | 0.16 | 0.33 ± 0.04 |
|  | C | 0.39 | 0.48 | 125.20 | 0 | 1 | 0.07 | 0.16 | 0.30 ± 0.04 |
|  | D | 0.50 | 0.50 | 100.02 | 0 | 1 | 0.03 | 0.22 | 0.12 ± 0.03 |
| TD<br>(Days) | A | 64.64 | 14.75 | 22.82 | 1 | 76 | 48.64 | 171.16 | 0.22 ± 0.04 |
|  | B | 50.82 | 10.74 | 21.14 | 23 | 63 | 39.69 | 79.38 | 0.33 ± 0.04 |
|  | C | 53.49 | 11.91 | 22.27 | 36 | 67 | 63.08 | 79.31 | 0.44 ± 0.04 |
|  | D | 39.43 | 7.93 | 20.13 | 26 | 60 | 12.28 | 50.9 | 0.19 ± 0.03 |

A total of 167,751 and 393,157 SNPs resulted in r² greater than 0.8 in 200K and 930K cross validation tests, respectively. Joining the SNPs originally genotyped and imputed it was possible to achieve approximately 600K SNPs imputed with high accuracy. **Supplementary Table 1** shows the number of SNPs before quality control and the number of SNPs used for meta-analysis post quality control for each population used in this study. After MAF and HWE filtering (see methods) of the imputed panel at 930K, the minimum number of markers passed the quality control range from 413,198 in POP D to 462,637 in POP A. This number of markers were used to estimate variance components, heritabilities and to perform GWAS analysis in each trait and challenge, which are also shown in Table 2. Significant SNP-heritabilities were estimated for both trait definitions in all challenges, ranging from 0.12 (POP D) to 0.33 (POP B) in BS and from 0.19 (POP D) to 0.44 (POP C) in TD.

### GWAS analyses

The results visualized as Manhattan plots of the GWAS of each challenge are available in the supplementary figures (Fig.S1-S4). Briefly, we found 2 SNPs associated with resistance as BS in POP B in *Ssa02*, whereas for POP A, C and D no associated SNPs were found. For TD, we found 1 SNP genome-wide significant SNP on *Ssa17* in POP A, 6 genome-wide significant SNPs on *Ssa02* in POP B, whereas no significantly associated SNPs were found in POP A, C nor D.

### Variant-based meta-analysis

The meta-analysis was performed using a total of 5,839 individuals and the output resulted in a total of 537,071 non-overlapping SNPs. Figure 1 shows the GWAS meta-analysis Manhattan plot for resistance to *P. salmonis* measured as BS and TD. A polygenic architecture was revealed by METAL results for both traits. We identified four genomic regions associated with resistance as BS. We found a total of 25 SNPs in these regions which were located on *Ssa02* (10), *Ssa11* (1), *Ssa24* (12) and *Ssa26* (2). For TD, we identified 177 SNPs located on *Ssa02* (165), *Ssa13* (1), *Ssa17* (7), *Ssa22* (1) and *Ssa29* (3). The Pearson correlation between the magnitude of the effect of markers on BS and TD was 0.58, indicating a moderate to high correlation between the genetic architecture of both traits (*p*-value < 2.2 e-16). However, only 1 SNP (*Ssa02* - 22420933) was identified as significant for both BS and TD: showed a negative effect direction (slope) in 3 of the 4 populations (removed from POP D during quality control (HWE)). This marker had a MAF of 0.18, 0.18 and 0.44 for POP A, B and C and explained from 0.5 to 1.1% of phenotypic variance (PVE) for BS and from 0.5 to 1.3% for TD among all populations, following the Teslovich et al. 2010 formula (Teslovich et al., 2010) (**Supplementary Table 2**).

**Figure 1.**
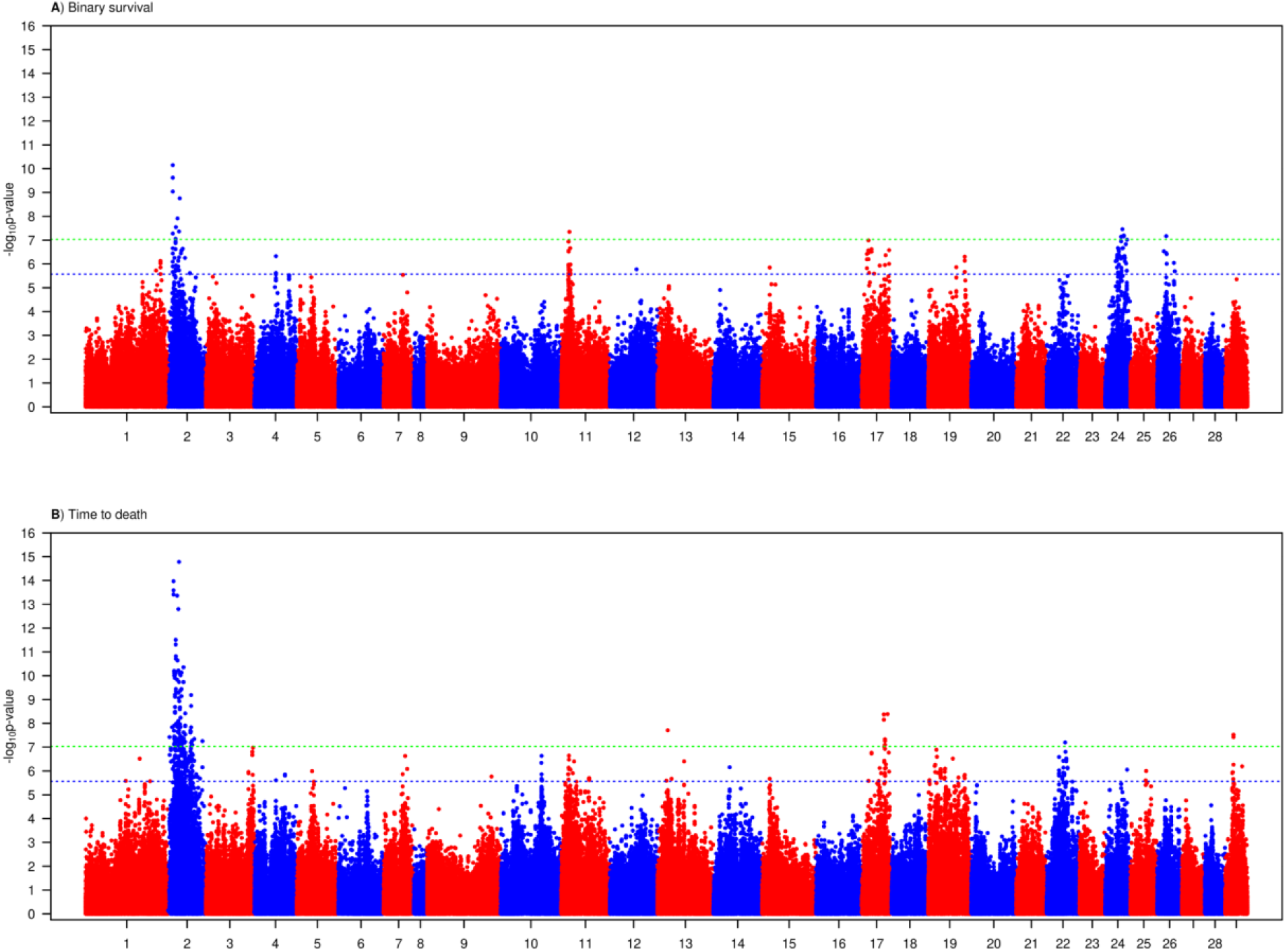
Manhattan plots of GWAS meta-analysis results using METAL of A) Binary survival and B) Time to death. Green line shows the genome-level significance whereas the blue line indicates the chromosome-level significance.

### Gene-based meta-analysis and enrichment

All variants present in each input were annotated to genes using ICSASG_v2 assembly, resulting in 33,365 genes that contained at least one variant SNP. A total of 24 and 40 genes were found to be associated with BS and TD, respectively (Figure 2). Among these, 16 genes were found to overlap between the two independently performed analyses, all of them on *Ssa02*, which includes a total of 122 SNPs within these genes (Table 3). These are the strongest candidate genes associated with resistance to *P. salmonis* in *Salmo salar*. **Supplementary Table 3 and 4** includes the complete list of genes associated with BS and TD, respectively, their *p*-values and *z*-scores. Briefly, we found *Tripartite motif-containing 33-like* (*TRIM33L*), *t-SNARE domain-containing protein 1-like* (*TSNARE1*), *Zinc Finger Protein 385D* (*ZNF385D*), *cathepsin K-like* (*CTSK*), *Zinc and Ring Finger Protein 2* (*ZNRF2*), *DC-STAMP Domain Containing 2* (*DCST2*), *SMAD Family Member 4* (*SMAD4*), *BMS1 Ribosome Biogenesis Factor* (*BMS1*), *TBC1 Domain Family Member 15* (*TBCD15*), *Leukocyte Receptor Cluster Member 8* (*LENG8*), *Mediator Of DNA Damage Checkpoint 1* (*MDC1*) and 4 uncharacterized loci (*LOC106580132, LOC106577156, LOC106577146* and *LOC106578317*).

**Figure 2.**
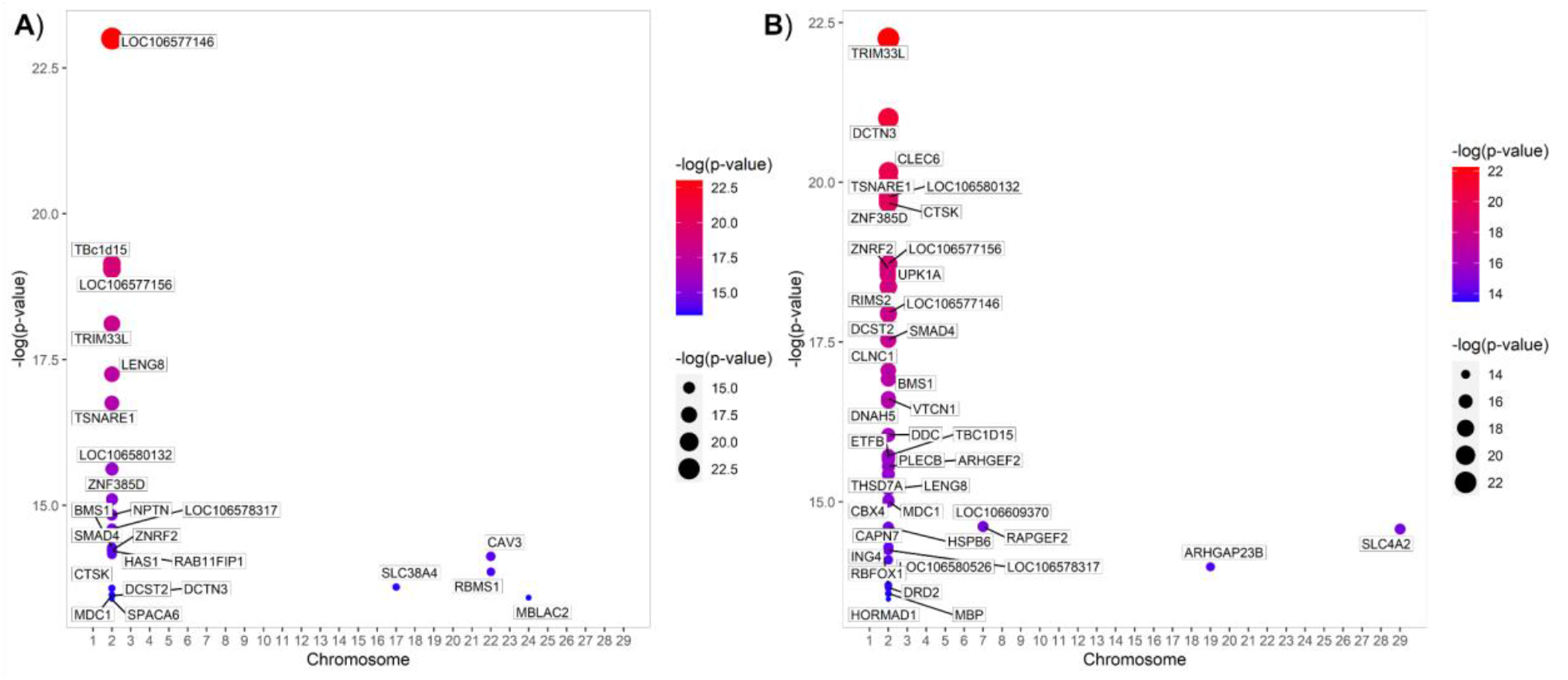
Gene analyses based on GWAS meta-analysis of A) Binary survival and B) Time to death. The x-axis represents the chromosome number, y-axis, dot color and size shows the negative log10-transformed gene-based *p*-value.

**Table 3.**
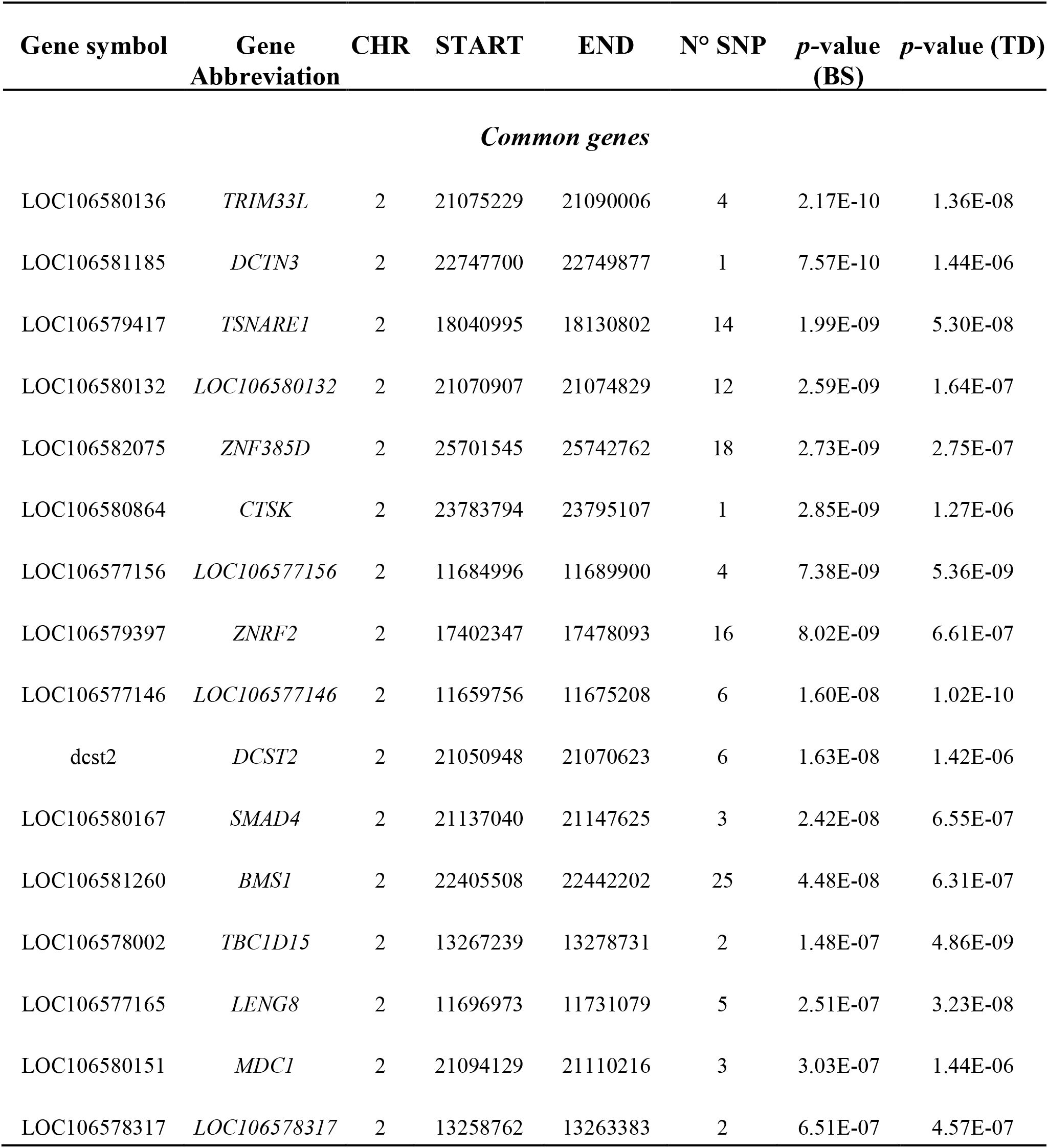
Common genes between BS and TD found by MAGMA. All genes p-values were below Bonferroni corrected threshold (*p*-value < 1.4 × 10^−6^ (=0.05/33,365)). Columns show gene ID, the chromosome number, start and end position for each gene, the number of SNPs included in each gene and their *p*-values for each trait.

Most of these genes are related in some way to molecular mechanisms of immune response, ribosome synthesis or homeostasis maintenance. Enrichment analysis revealed biological processes such as transcriptional regulation, ribosomal biogenesis, cell division, lipid storage, the transforming growth factor-beta signaling cascade and intracellular protein transport, cell adhesion, among others, for BS. On the other hand, for TD, the processes of greatest association with the genes found were transmembrane transport, signal transduction, proteolysis, transcription regulation, cell shape regulation, ribosome biogenesis and spermatogenesis, among others. Using the genes found in common for both traits we found that the most enriched processes were transcription regulation, cell shape regulation, ribosome biogenesis, intracellular protein transport and transforming growth factor beta receptor signaling pathway. **Supplementary File 1** contains genes associated with GO terms, their GO parent terms for BS, TD and for the common genes.

## DISCUSSION

This work was motivated out of a wish to better elucidate the genetic architecture of resistance to *P. salmonis* but also to detect with greater statistical power which genes might be associated with these traits. In this study we used the two trait definitions of resistance to *P. salmonis* most used throughout the literature and performed meta-analyses to reveal in detail the genetic architecture of both traits using METAL but also to detect candidate genes for both traits using MAGMA. At the center of the discussion are markers and genes that overlap between both trait meta-analyses and are discussed below.

### Genetic architecture of *P. salmonis* resistance

The results of this work supports a polygenic architecture of SRS genetic resistance as shown by previous studies (Correa et al., 2015; Moraleda et al., 2021; Yáñez et al., 2019). We found highly significant QTL on *Ssa*02 and *Ssa*24 for BS and *Ssa*02, *Ssa*17 and *Ssa*21 for TD. The SNPs found in this study are novel markers associated with SRS since they have not been associated with genetic resistance to *P. salmonis* in previous studies. We found a moderate correlation between the SNP effect (0.58), indicating that although there is a degree of correlation between BS and TD genetic architecture, there is a non-common component that differentiates the two traits and may explain the differences in the number of common SNPs found between these traits.

The polygenic nature of SRS-resistance is taken into account by the breeding company which the data in this study originate from. Currently, SRS-resistance is selected for using genomic selection, as are all other traits except those with strictly monogenic inheritance. Earlier, however, marker-assisted selection was used. Based on large-scale data from the field, this marker-assisted selection for SRS-resistance was shown to contribute effectively to reducing mortality from SRS (Evalualición de Variables de Cultivo sobre la Presentación de SRS y Use de Antibiótics, Aquabench report, May 2022).

The architecture of SRS resistance has been previously evaluated in Atlantic salmon in different studies (Correa et al., 2015; Moraleda et al., 2021; Yáñez et al., 2019). Yañez et al. 2019 found 3 SNP associated with BS distributed among Ssa01 and Ssa21, and 10 SNP associated with TD distributed on Ssa01, Ssa03, Ssa12, Ssa16, Ssa17 and Ssa22, with no overlapping SNPs between traits. The study considered a single population, a 50K SNP density, and the markers found are not the same as those found in the present work. Moraleda et al. 2021 found 6 SNPs associated with BS distributed on Ssa01, Ssa02, Ssa12 and Ssa27. This study used a single population, imputed from 1K to 38K SNP to perform the GWA. None of these significant SNPs are common with the present work’s significant SNP. Correa et al. 2015 found 2 common significant SNP for BS and TD on Ssa01, and 3 non-common within Ssa01 and Ssa17. None of the above were found on the present work. However, we found 7 SNPs highly significant in *Ssa17* for TD, where the closer one is located less than 901 kB away from that one found by Correa et al. 2015. Markers linked to potential causal or major effect markers may appear as significant in the GWAS, together with the effect of the number of markers used to test their effect and define significance.

Differences in genetic architecture between SRS resistance definitions have been observed in previous studies in other species such as coho salmon (Barría et al., 2018) and rainbow trout (Barria et al., 2019). Therefore, finding differences in the genetic architecture of both definitions of genetic resistance is not a novel phenomenon.

In this meta-analysis, one QTL on chromosome Ssa02 stood out as much more significant than other QTLs. However, this QTL was not prominent in earlier studies. There could be several reasons behind this finding. First, the studies were performed on genetically distinct populations of Atlantic salmon; although all or most Atlantic salmon populations present in Chile ultimately originate from Norway, the different populations are known to originate from only partly overlapping sets of rivers. Differences in experimental setup could also lead to different results across studies. For instance, the experimental challenge conditions (Intraperitoneal (IP) vs. Cohabitation (COH)), the statistical model, softwares, the number of samples and the number of SNPs and their origin (different SNP-chips, imputed versus genotyped SNPs) may influence the results. The pattern of infection of IP differs from that of COH infections (Almendras et al., 1997) and IP is not a natural form of infection, as the main route of entry for *P. salmonis* is via epithelial tissues (skin and gills) (Smith et al., 1999). Epidermal mucus and epithelial tissues are a major determinant and the main barrier against bacterial infections (Dash et al., 2018) and IP challenges bypass this, leading to a partial capture of genetic resistance being assessed. The 4 challenges used in our study were developed using a cohabitation scheme, assimilating the natural route of infection. Furthermore, unlike the earlier studies where LF-89 strain were used, the present work uses EM-90 strain, which has a higher host preference in Atlantic Salmon than in other farmed salmonid species and shows higher mortalities and tissue damage compared to LF-89 (Rozas-Serri et al., 2017; Saavedra et al., 2017). Pan-genomic studies have shown differences in strain-specific virulence factors related to adherence, colonization and endotoxins, which may directly be associated in differences in pathogenesis and host-pathogen interactions (Nourdin-Galindo et al., 2017; Rozas-Serri et al., 2018), however it is unknown whether and to what extent strain influences the genetic architecture of genetic resistance. We hypothesize that the combinatorial of the factors mentioned above could influence mortality and consequently phenotypes that are evaluated, affecting the detection of QTLs within and between studies.

The proportion of the phenotypic variance explained by the most significant markers found by Correa et al. 2015 did not exceed 0.45%, while the common SNP for both traits found in this study explains up to 1.3%. This marker was imputed in all populations, as it is originally a 930K panel SNP. The minor allele frequency of this marker is intermediate (POP A and B) to high (POP C) as shown in **Supplementary Table 2**.

### Candidate genes and biological processes associated with both definitions of *P. salmonis* resistance

The gene-based meta-analysis detected a total of 16 genes that are shared between both definitions of genetic resistance to *P. salmonis* (Table 3). We found *SMAD4*, a transcription factor induced by TGF-β signaling that affects muscle regeneration and myogenesis (Lee et al., 2005). This gene was associated with two GO terms: via transforming growth factor (GO:0007179) and regulation of transcription, DNA-templated (GO:0006355). *SMAD4* is a gene whose transcripts are induced via transforming growth factor (TGF-β), the latter being a generally anti-inflammatory cytokine secreted by platelets and macrophages following injury, where it attracts neutrophils, macrophages and fibroblasts to confront the causative agents of the injury (parasites, viruses, bacterias or others) (Lilleeng et al., 2009; Maehr et al., 2013; Skugor et al., 2008; Werner & Grose, 2003; M. Yang et al., 2012). Interestingly, SMAD4 has also been associated with transcriptomic profiling in response to *P. salmonis* in Atlantic salmon (Tacchi et al., 2011) and proteomic profiling in salmon macrophage-like cells (Ortiz-Severín et al., 2020).

Among the genes with the highest number of associated SNPs we found *ZNRF2* (*E3 ubiquitin-protein ligase ZNRF2*) with 16 SNPs associated and with three GO terms: protein ubiquitination (GO:0070936), proteasome-mediated ubiquitin-dependent protein catabolic process (GO:0043161) and with protein K48-linked ubiquitination (GO:0070936). This gene belongs to a gene family that are important biomarkers in the ubiquitin–proteasome system, one of the main protein degradation pathways in living organisms (Cai et al., 2022). This gene is important for protein translation and antigen processing (Gao & Karin, 2005) and has been found to be down-regulated in spleen following SRS infections (Valenzuela-Miranda & Gallardo-Escárate, 2018).

We found MDC1 (Mediator of DNA damage checkpoint protein 1) associated with three GO terms: lipid storage (GO:0019915), pigment accumulation (GO:0043476) and with positive regulation of sequestering of triglyceride (GO:0010890). This gene has been related to DNA repair via recruiting DNA damage response factors as well as in regulation of mitosis (Eliezer et al., 2009; Leimbacher et al., 2019). We also found genes such as *TSNARE1* associated with intracellular protein transport (GO:0006886) and vesicle-mediated transport (GO:0016192). This gene is highly associated with intracellular protein transport endosomal trafficking (Plooster et al., 2021). Finally, we also found the *BMS1* gene (*ribosome biogenesis protein BMS1*), with 25 associated SNPs and one GO term associated (ribosome biogenesis: GO:0042254). Initiates ribosome biogenesis, regulates early rRNA processing and it is involved in ribosome biogenesis, translation capacity and protein folding. Infection with SRS can interfere with host protein synthesis and suppress the innate immune response via CsrA superfamily virulence-related proteins which are RNA-binding proteins and global regulators of carbohydrate metabolism genes facilitating mRNA decay (Barandun et al., 2018; Wang et al., 2012). Rapid responsiveness at mRNA level is imperative to establish an immune response to infection. *P. salmonis* alters cytoskeleton remodeling, intracellular transport, organelle organization, vesicle and endosomal trafficking, inhibition of the antioxidant response (Rozas-Serri, 2022).

On the other hand, we found genes previously described to be differentially expressed in different studies. For example, *LOC106577156, LOC106578317, LOC10658013* and *TBC1D15* were found by Moraleda et al. 2021. *TBC1D15* was found up-regulated in transcriptomic profiles of Atlantic salmon with high and low loads of *P. salmonis* (Xue et al., 2021). An important gene we found was *Tripartite motif-containing 33-like* (*TRIM33L*), a gene that codes for a nuclear protein that associates with specific DNA-binding transcription factors to modulate gene expression. This gene has been cataloged as an essential for macrophage and neutrophil mobilization in response to inflammatory recruitment signals, including bacterial infections (Demy et al., 2017) and viremias (Kim et al., 2020). *TRIM33L* was previously detected in a wssGWAS analysis for economically-important traits in rainbow trout (Gonzalez-Pena et al., 2016). This gene appears to have a function in maintaining myoblast populations through myogenic precursor cell proliferation, and is up-regulated during muscle regeneration in model species (Mohassel et al., 2015) and rainbow-trout (Gonzalez-Pena et al., 2016).

The genes mentioned in this section are the most statistically significant genes and potentially play a functional role in the genetic resistance to SRS in Atlantic salmon. Not only because they have been detected in the present work with a statistical power unprecedented in the literature but also because some of them have been previously associated with SRS. The results of this work could guide functional genomics studies to understand in greater depth the role of these genes on the evaluated phenotypes.

## CONCLUSIONS

This manuscript presents the first variant-level meta-analysis to describe the genetic architecture as well as the first gene-level meta-analysis to present candidate genes associated with resistance to *Piscirickettsia salmonis*. The candidate genes detected in this work could form a baseline for functional studies or gene editing. Additionally, these markers could be used for genomic prediction studies in order to prioritize markers for a cost-effective strategy to select resistant animals against this disease.

## ACKNOWLEDGMENTS

We thank AquaGen Chile and Norway for providing the data and for their close collaboration. This research was carried out under the collaborative project between the Aquaculture Genomics Laboratory of the University of Chile and AquaGen.

